# Microbial loop of a Proterozoic ocean analogue

**DOI:** 10.1101/2021.08.17.456685

**Authors:** Jaspreet S Saini, Christel Hassler, Rachel Cable, Marion Fourquez, Francesco Danza, Samuele Roman, Mauro Tonolla, Nicola Storelli, Stéphan Jacquet, Evgeny M. Zdobnov, Melissa B. Duhaime

**Affiliations:** Department F.-A Forel for Environmental and Aquatic Sciences, Earth and Environmental Sciences, University of Geneva, Switzerland; Department of Genetic Medicine and Development, University of Geneva, and Swiss Institute of Bioinformatics, Geneva, Switzerland; Department of Ecology and Evolutionary Biology, University of Michigan, Ann Arbor, MI, United States; Institute of Microbiology (IM), Department for environmental constructions and design (DACD), University of Applied Sciences and Arts of Southern Switzerland (SUPSI), Switzerland; Department of Botany and Plant Biology, University of Geneva, Switzerland; Aix Marseille Univ., Université de Toulon, CNRS, IRD, MIO UM 110, 13288, Marseille, France; Université Savoie Mont-Blanc, INRAE, UMR CARRTEL, Thonon-les-Bains, France

## Abstract

Meromictic Lake Cadagno, an ancient ocean analogue, is known for its permanent stratification and persistent anoxygenic microbial bloom within the chemocline. Although the anaerobic microbial ecology of the lake has been extensively studied for at least 25 years, a comprehensive picture of the microbial food web linking the bacterial layer to phytoplankton and viruses, with explicit measures of primary and secondary production, is still missing. This study sought to understand better the abundances and productivity of microbes in the context of nutrient biogeochemical cycling across the stratified zones of Lake Cadagno. Photosynthetic pigments and chloroplast 16S rRNA gene phylogenies suggested the presence of eukaryotic phytoplankton through the water column. Evidence supported high abundances of *Ankyra judayi*, a high-alpine adapted chlorophyte, in the oxic mixolimnion where oxygenic-primary production peaked. Through the low- and no-oxygen chemocline and monimolimnion, chlorophytes related to *Closteriopsis acicularis*, a known genus of meromictic lakes, and *Parachlorella kessleri* were observed. *Chromatium*, anoxygenic phototrophic sulfur bacteria, dominated the chemocline along with *Lentimicrobium*, a genus of known fermenters whose abundance was newly reported in Lake Cadagno. Secondary production peaked in the chemocline suggesting primary producers depend on heterotrophs for nutrient remineralization. As previously observed, sulfur-reducing bacteria (SRBs), especially *Desulfocapsa* and *Desulfobulbus*, were present in the chemocline and anoxic monimolimnion. Virus-to-microbe ratios (VMR) peaked in the zone of phytoplankton yet were at a minimum at the peak of *Chromatium*. These dynamic trends suggest viruses may play a role in the modulation of oxygenic and anoxygenic photo- and chemosynthesis in Lake Cadagno and other permanently stratified systems.

**Importance:** As a window to the past, the study offers insights into the role of microbial guilds of Proterozoic ocean chemoclines in the production and recycling of organic matter of sulfur- and ammonia-containing ancient oceans. The new observations described here suggest that eukaryotic algae were persistent in the low oxygen upper-chemocline in association with purple and green sulfur bacteria in the lower half of the chemocline. Further, this study provides the first insights into Lake Cadagno viral ecology. High viral abundances suggested viruses may be essential components of the chemocline where their activity may result in the release and recycling of organic matter. The framework developed in this study through the integration of diverse geochemical and biological data types lays the foundation for future studies to quantitatively resolve the processes performed by discrete populations comprising the microbial loop in this early anoxic ocean analogue.

## Introduction

Roughly 2.3 billion years (Gyr) ago the oceans were filled with sulfur ^1^ and anoxygenic photosynthesis was a dominant mode of microbial primary production ^2–4^. Following the great oxidation event ∼2 Gyr, anoxygenic photosynthesis continued to sustain microbial life in the euxinic environments below oxygenated surface waters ^1,3,5^. Habitats in which anoxygenic photosynthesis significantly contributes to primary production continue to persist, such as in anoxic zones of permanently stratified sulfur-rich meromictic lakes, like Lake Cadagno ^6^ where hydrogen sulfide serves as an important electron donor ^7^. These permanently stratified lakes are resistant to seasonal mixing events that would otherwise redistribute oxygen throughout the water column ^8^. In these systems, biogeochemical processes provide a link between the contrasting dominant modes of primary production, thereby directly connecting the food webs of the oxygenated and anoxic habitats.

The microbial community composition of meromictic Lake Cadagno typifies the reconstructed signatures of life in the Proterozoic ocean ^9,10^. Purple and green sulfur bacteria (PSB and GSB, respectively) are amongst the hallmark bloom-forming microbes that are frequently observed in the chemocline of Lake Cadagno ^11,12^. Within the chemocline, carbon, nitrogen, sulfur, and iron cycling are integrated through metabolic activities of the anoxygenic photosynthetic purple sulfur and green sulfur bacteria ^13–16^. Two PSB, *Thiodictyon syntrophicum* and *Chromatium okenii*, have been identified as the greatest contributors to primary production in Lake Cadagno ^12,14,17^. However, recent measurements of primary production to date have been limited to the chemocline ^12,18^ and measurements of secondary production do not exist. Phytoplankton (including modern eukaryotes and cyanobacteria) likely coexisted with anoxygenic phototrophic sulfur bacteria in Proterozoic oceans ^19,20^. High phytoplankton abundances and productivity were expected in the oxic zone because phytoplankton produces organic matter via oxygenic photosynthesis. In order to better understand microbial loop linkages across the stratified mixolimnion-chemocline transition in Lake Cadagno, paired measurements of productivity, microbial community composition, and diversity are essential next steps.

Lake Cadagno studies to date have focused on the phytoplankton and prokaryotic members of the microbial community and have yet to include viruses. Viruses are known to impact both phytoplankton and heterotrophic microbial populations through lysis and lysogeny and may hold a central position in the marine food webs and biogeochemical cycling ^21,22^. Through the lysis of their microbial hosts, viruses play a role in the conversion of particulate organic carbon to dissolved organic carbon, which is predicted to be capable of sustaining the microbial loop ^23,24^. Viruses have been identified in meromictic lakes ^25–27^, but their abundances across the vertical water column of Lake Cadagno are unknown.

To better understand the role the microbial loop plays in sustaining the food web across the stratified layers of Lake Cadagno, microbial community members (phytoplankton, prokaryotes, and viruses) were studied in the context of biogeochemical and primary and secondary production measurements across the oxic-anoxic transitions of the meromictic Lake Cadagno. We hypothesized the mixolimnion to have low primary production relative to the chemocline because the chemocline is known to be inhabited by a persistent microbial bloom of primary producing phototrophic sulfur bacteria ^12^. We also expected high secondary production and viral abundance to be an indicator of effective organic matter recycling within the Lake Cadagno water column, because photoautotrophs, both phototrophic sulfur bacteria and phytoplankton, rely on heterotrophs and viruses for the remineralization of organic matter and nutrient cycling ^28,29^. Through a combination of physical, chemical and biological analyses, this work provides new evidence on how transitions between permanently stratified lake habitats and assemblages may sustain microbial food webs, informing our understanding of the aquatic ecosystems of early Earth.

## Results

### Position of Mixolimnion, Chemocline, and Monimolimnion

Transitions between the major stratified layers sampled from Lake Cadagno were visually apparent by the colour of pigmented biomass captured on the 0.22 µm filters (Fig. 1A). From the surface to 12.5 m depth, the oxygen levels fell from a maximum of 9.24 mg/L to 2.79 mg/L (light green, day 2; Fig. 1B). As hypoxia is defined as 2.00 mg O_2_/L ^30^, 12.5 m marks the boundary of the oxic mixolimnion zone and the upper chemocline (top light purple; Fig. 1). The chemocline was 13.5-15.5 m, as determined based on near-zero oxygen and light levels (day 2), peaked turbidity, a slight decrease in temperature, and a simultaneous rise in conductivity (dark purple; Fig. 1, B-D). At 14.5 m, oxygen levels were below detection and <1% of surface light penetrated (day 2; Fig. 1B, D). Within the chemocline, the deep chlorophyll maximum (DCM) and peak turbidity shifted overnight between days 1 and 2, but in both cases, the peak turbidity was observed 0.5-1.5 m below the DCM (day 2; Fig. 1B, D). Peak phycocyanin concentrations (26.53 µg/L) and basal fluorescence (Fb = 66.0) were observed at the DCM (day 2, 13.5-14 m) and above a secondary chlorophyll peak at 15 m (Fig. 1D, Fig. S1E). The monimolimnion was defined as the zone between 15.5 m and the benthos. The peak in particulate sulfur (0.74 ppm), H_2_S (2.84 mg/L) and ammonia (262.4 µg/L) concentrations were observed in the monimolimnion, as well as a decline in turbidity and photosynthetic pigments (Fig. 1B, E, H).

**Figure 1.**
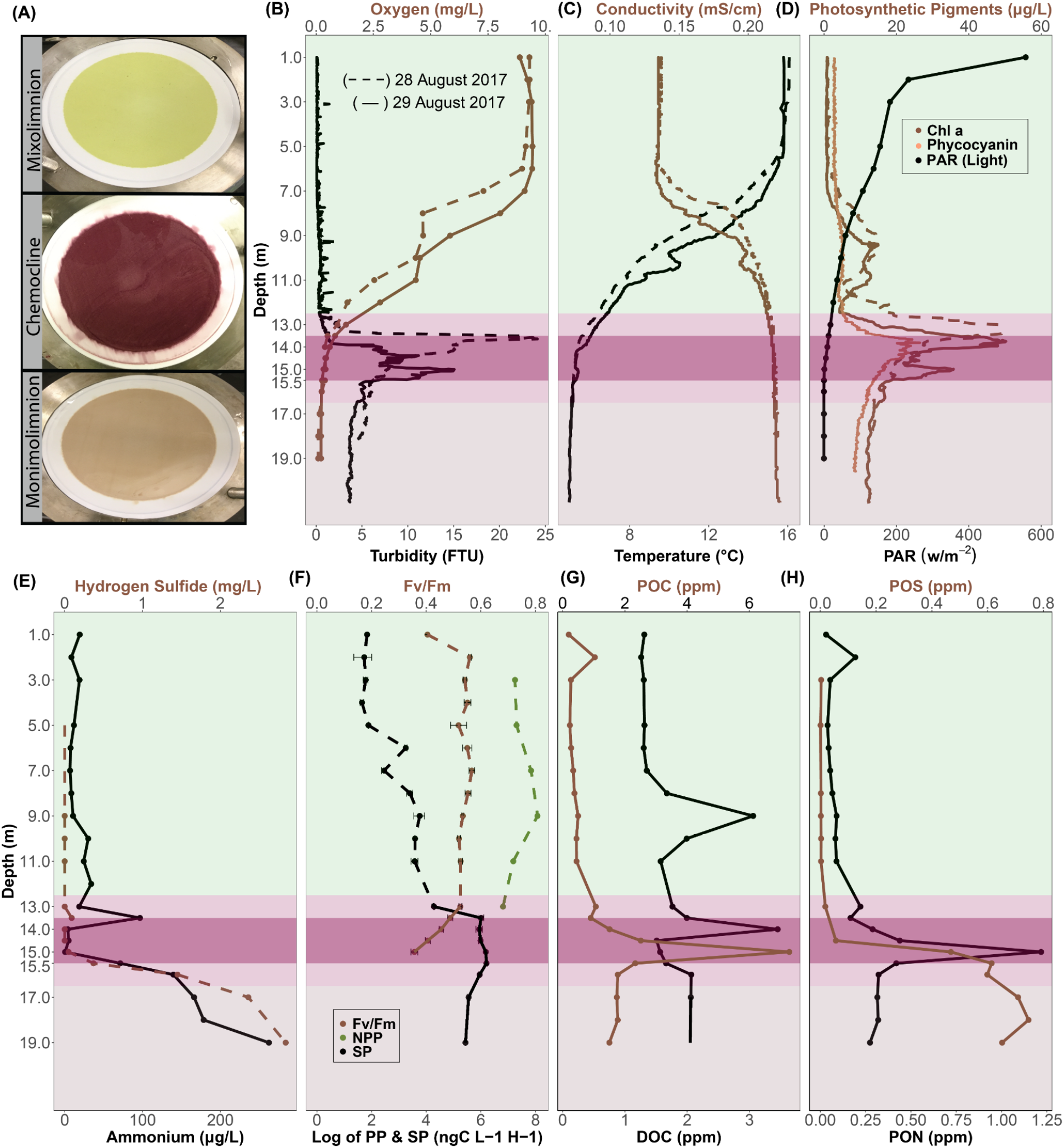
Depth profiles of biogeochemical parameters of Lake Cadagno. (A) Photographs of biomass collected on 0.22 µm (142 mm diameter) filters from Lake Cadagno mixolimnion, chemocline, and monimolimnion strata. The depth profiles of (B) oxygen and turbidity, (C) conductivity and temperature, (D) Chl *a*, phycocyanin, and photosynthetically active radiation (PAR) (E) hydrogen sulfide and ammonium concentrations, (F) net primary production (NPP) and secondary production (SP) rates, (G) particulate organic carbon (POC) and dissolved organic carbon (DOC) concentrations, (F) particulate organic sulfur (POS) and particulate organic nitrogen (PON) concentrations. In B-H, the mixolimnion, chemocline transition zone, chemocline, and monimolimnion layers are indicated by light green, lilac, purple and light brown backgrounds, respectively. Dashed and solid lines represent 28th and 29th August 2017, respectively.

### Microbial production at the oxic/anoxic interface

Between 7.0-11.0 m in the lower mixolimnion, there was a slight rise (up to 13.12 µg/L on day 1 and 11.80 µg/L on day 2) in Chl *a* (Fig. 1D). The Chl *a* peak in the mixolimnion at 9.0 m corresponded with a peak in net primary production (NPP; 3210 ng C L^-1^ h^-1^; Fig. 1D, F). The NPP peak declined in the lower mixolimnion (11.0 m) and upper chemocline (13.0 m). Though NPP was not measured below 13.0 m, basal and maximum fluorescence measured through the chemocline gave an indication of phytoplankton photosynthetic activity and efficiency (Fig. S1E). In the oxic-anoxic boundary, a maximum Fv/Fm (an indicator of photosynthetic efficiency) of 0.49 was observed between 13.0-13.5 m and declined to 0.36 at 15.0 m (Fig. 1F) where the light was absent. Overlapping with this oxygenic photosynthesis, the highest rates of secondary productivity (SP) were observed through the chemocline (Fig.1F). SP was significantly positively associated with some indicators of both photosynthetic microbes (Pearson: Chl *a*: R=0.96, p-value=1.7e-5; phycocyanin: R=0.80, p-value=0.005) and total biomass (Pearson: turbidity: R=0.78, p-value=0.007; total cell counts: R=0.65, p-value=0.04) (Fig. S2). In the chemocline, SP and indicators of NPP (phycocyanin, Chl *a*) were significantly negatively correlated with oxygen and light (Pearson: R=-0.7 to -0.9, p-values = 0.0002 to 0.01; Fig.S2). The peak of DOC (14.0 m) was followed by peaks of POC, PON (15.0 m; Fig. 1G-H). In the monimolimnion below the chemocline, some organic and inorganic compounds continued to rise (POS, H_2_S, NH_4_^+^), while PON and POC dropped (Fig. 1).

### Prokaryote- and Virus-like-particle abundances and trends

The concentration of prokaryotic-like particles (PLPs; prokaryotic cell counts inferred from flow cytometry data) was highest in the chemocline with a peak of 905,000 cells/ml ± 28,583 at 15.0 m (496,000 ± 17,295 avg cells/ml), roughly half that in the monimolimnion (435,833 ± 10,243 avg cells/ml), and was lowest in the mixolimnion (109,556 ± 3,496 avg cells/ml). In contrast, virus-like particles (VLPs; viral counts inferred from flow cytometry data) were highest at the bottom of the mixolimnion (1.71e+08 ± 2.63e+07 VLP/ml at 11.0 m), lowest in the upper mixolimnion (7.03e+07 ± 1.50e+07 VLP/ml at 5.0 m), and relatively invariable through the rest of the water column (Fig. 2B). As a result, the virus to microbe ratio (VMR) peaked with viruses at the bottom of the mixolimnion but decreased drastically when prokaryotic-like particles rose in the chemocline (Fig. 2C).

**Figure 2.**
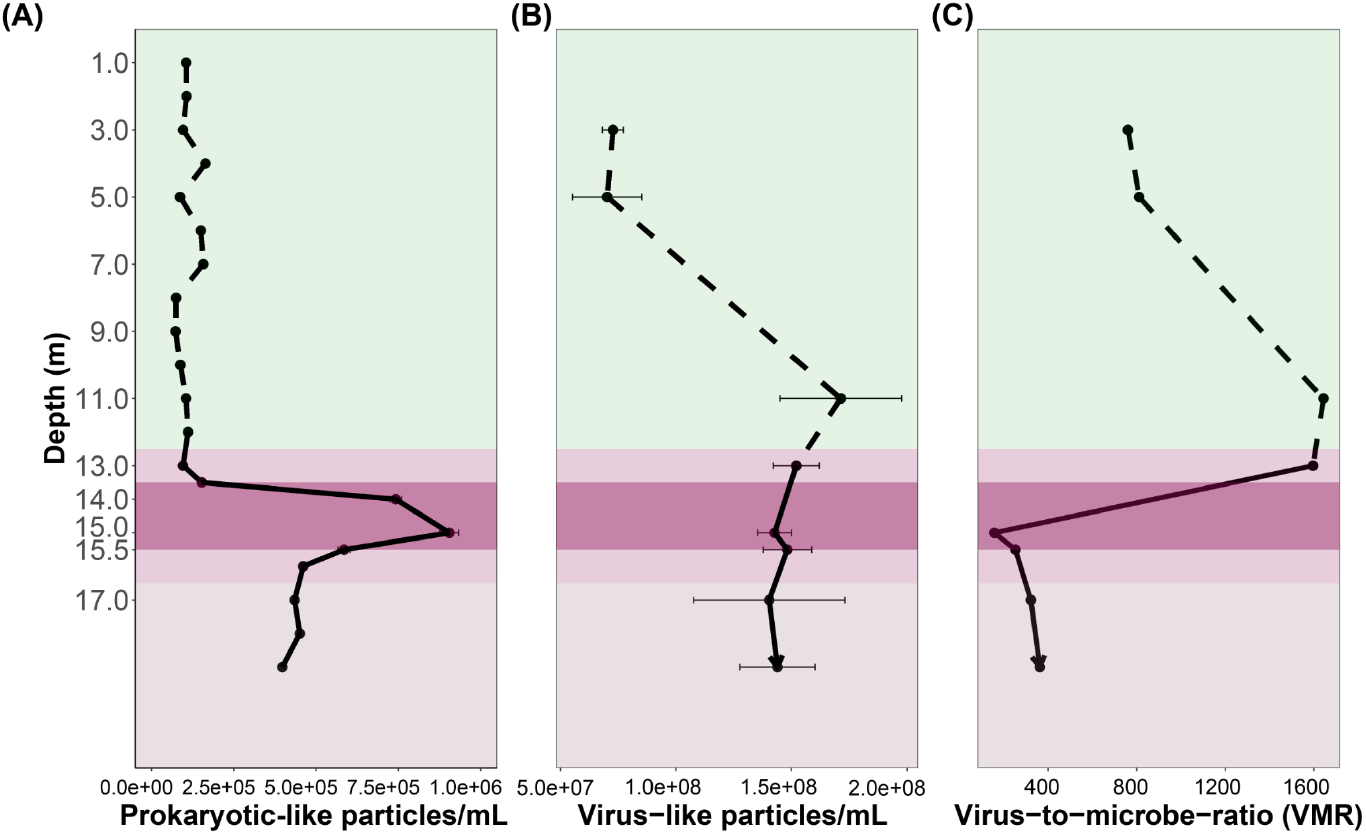
The abundance of (A) prokaryote-like particles (PLP) and (B) virus-like particles (VLP) from flow cytometry analyses and (C) virus-to-microbe ratio, as VLP/PLP.

Significant positive correlations were observed between PLP concentrations and other indicators of biomass: turbidity (R=0.95; p-value=0.00003), Chl *a* (R=0.76; p-value=0.01), POC R=0.92; p-value=0.0001), PON (R=0.93; p-value=0.0001), and POS R=0.76; p-value=0.010) (Fig. S3). Of those, only the VLP concentrations were correlated with Chl *a* (R=0.77; p-value=0.02, (Fig. S4). Significant negative correlations were observed between VLPs and oxygen (R=-0.82, p-value=0.01) and light R=-0.90, 0.002). Significant positive correlations were observed between VLP and depth (R=0.78, p-value=0.02), conductivity (R=0.92, p-value=0.001), and SP (R=0.70; p-value=0.05) (Fig. S4).

### Bacterial community composition through the mixolimnion, chemocline, and monimolimnion

Both relative (OTU count scaled by total OTUs) and absolute abundances (OTU count scaled by total FCM PLP counts^31^) were considered to assess abundances of microbial taxa through Lake Cadagno (Fig, 3A-B). Absolute abundances of Bacteria-classified cells (‘Bacteria’ OTU count scaled by total FCM PLP counts^31^) were at a minimum in the oxic mixolimnion (day 1; 0-12.5 m; Fig. 3B, Table 1). On average, the mixolimnion community primarily consisted of Actinobacteria (35.34% of total OTUs; 33,815 cells/ml), Bacteroidetes (21.47% of total OTUs; 20,541 cells/ml), Proteobacteria (20.45% of total OTUs; 19,563 cells/ml), and Cyanobacteria (5.4% of total OTUs; 5,225 cells/ml) phyla (Fig. 3A-B, Fig. S5).

**Figure 3:**
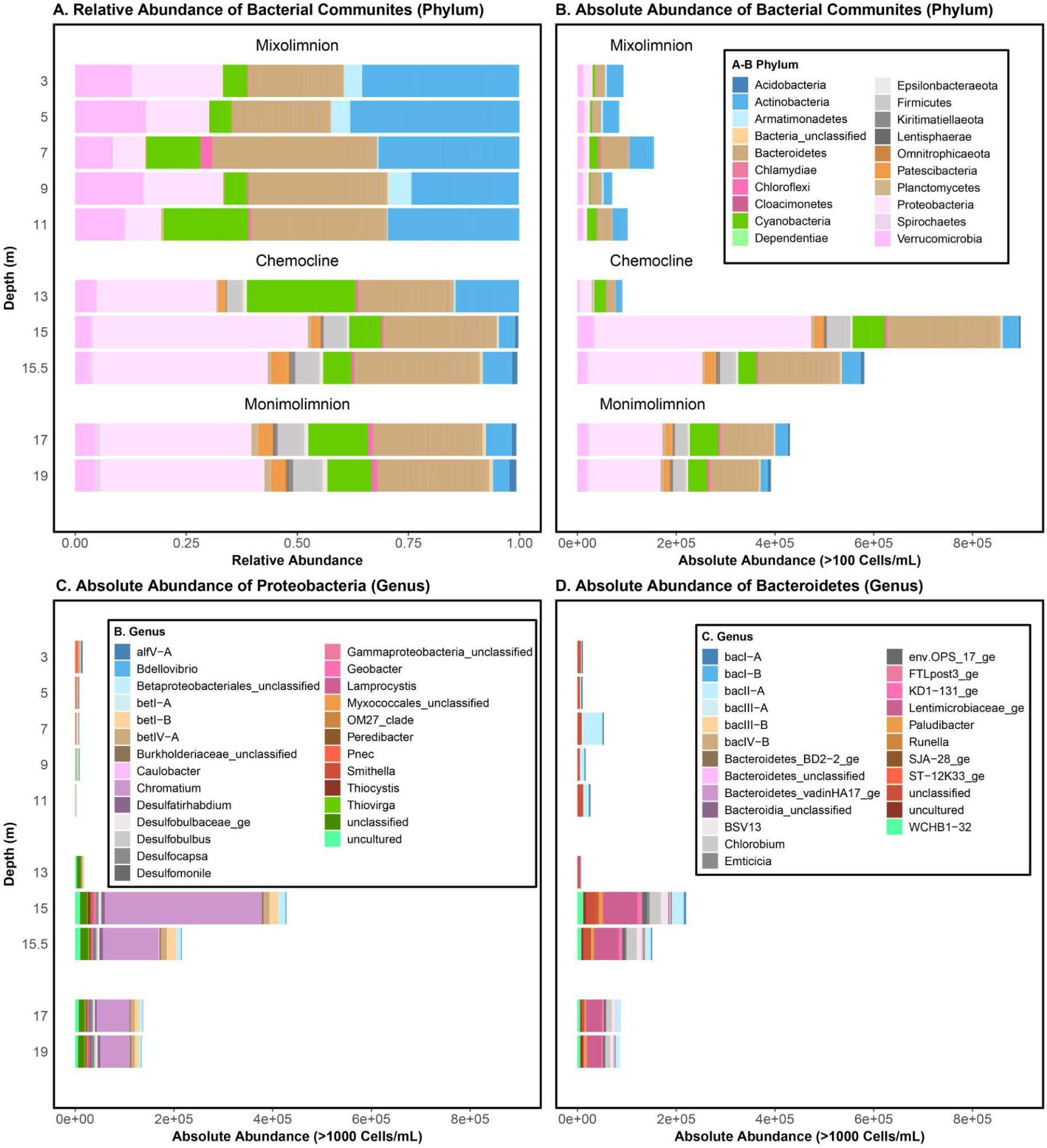
(A) Relative (OTU count scaled by total OTUs) and (B) absolute (OTU count scaled by total FCM PLP counts^31^) abundances of bacterial phyla at the sampled depths in Lake Cadagno, (C) genera in the Proteobacteria phylum, (D) genera in the Bacteroidetes phylum. To the right of each panel, the mixolimnion, chemocline transition zone, chemocline, and monimolimnion layers are indicated by light green, lilac, purple and light brown backgrounds, respectively and correspond with the depths reported on the left-most y-axis.

In the chemocline, the absolute abundance of Bacteria-classified cells peaked at 15 m (day 2; Fig. 3A), which corresponded with peak turbidity, POC and PON (Fig. 1). The major phyla represented at this depth were Proteobacteria (48.42% of total OTUs; 438,272 cells/ml) and Bacteroidetes (25.35% of total OTUs; 229,448 cells/ml). The genera that dominated these phyla at 15 m were purple sulfur bacteria (PSB) *Chromatium* (35.18% of total OTUs; 318,381 cells/ml). *Chromatium* (Proteobacteria) was significantly positively correlated with turbidity (R=0.93; p-value=7.3e-5) and total cell counts (R=0.99; p-value=8.2e-09) (Fig.6 A-B). An unclassified *Lentimicrobiaceae* OTU, the second most abundant OTU in the chemocline at 15 m, was significantly positively correlated with secondary production (*R=*0.8; p-value*=*0.005), turbidity (R=0.91, p-value=0.0002), PLPs (R=0.85, p-value=0.001), PON (R=0.71, p-value=0.02), POS (R=0.92; p-value=5e-04), yet, negatively correlated with light (*R=*-0.78, p-value*=*0.007, Fig.S6H) and oxygen concentration (*R=-*0.87; p-value*=*0.001; Fig. S6 C-I). Though Firmicutes was not a dominant phylum itself, an unclassified genus of the phylum, *Erysipelotrichaceae* (3.90% of total OTUs; 35,360 cells/ml), was among the dominant genera at 15 m (Table 1). Green sulfur bacteria (GSB), *Chlorobium* (2.29% of total OTUs, 20,808 cells/ml), was the fifth most abundant genera of chemocline. Previously observed phototrophic sulfur bacteria, *Thiodictyon* and *Lamprocystis*^*12,15*^, and sulfur-reducing *Desulfocapsa* and *Desulfobulbus* were present at less than 0.01% of total OTUs in the chemocline (Table 1).

On average, the total PLP counts in the monimolimnion (17.0-19.0 m) were less than half that of the peak chemocline counts and more than twice the average mixolimnion counts. The average relative and absolute (Fig. 3A-B) abundances of the dominant populations reflected those of the chemocline at 15 m, which included Proteobacteria (34.05% of total OTUs; 148,019 cells/ml), Bacteroidetes (24.53% of total OTUs; 106,635 cells/ml), Cyanobacteria (13.44% of total OTUs; 58,436 cells/ml), and Actinobacteria (5.8% of total OTUs; 25,301 cells/ml). In addition to bacteria, archaeal OTUs of the *Methanoregula* and *Woesearchaeia* genera were identified in the monimolimnion, but they were not further considered owing to their overall low abundances (<100 OTUs, Table 2).

### Oxygenic phototrophs in the mixolimnion, chemocline, and monimolimnion

Cyanobacteria was among the top phyla consistently observed at each depth in the water column, but at no depth did it dominate (Fig. 3A). At all depths, *Oxyphotobacteria* and chloroplast-classified OTUs dominated the Cyanobacteria, representing 94-100% of the total Cyanobacteria-classified OTUs (Fig. 4A). Of the low abundance cyanobacterial genera, *Cyanobium* was found throughout the water column, while *Pseudanabaena* and *Gastranaerophilales* were predominantly found at 15 m and deeper (Fig. 4A-B). To better understand the relationship between the microbial guilds of Lake Cadagno, we sought more evidence regarding the origin of these putative chloroplast OTUs. The phylogeny of all Cyanobacteria-classified OTUs and their two nearest neighbours in SILVA ^32^ indicated that the chloroplast and *Oxyphotobacteria* OTUs clustered in clades with chloroplast 16S rRNA genes of cultured eukaryotic algae (Fig.4B): Chlorophyta (Otu00008, Otu000342, Otu00006, Otu00323), Ochrophyta (Otu00033), Streptophyta (Otu00208), Haptophyta (Otu00143), and uncultured-chloroplasts (Otu00042, Otu00051, Otu00052, Otu00055).

**Figure. 4:**
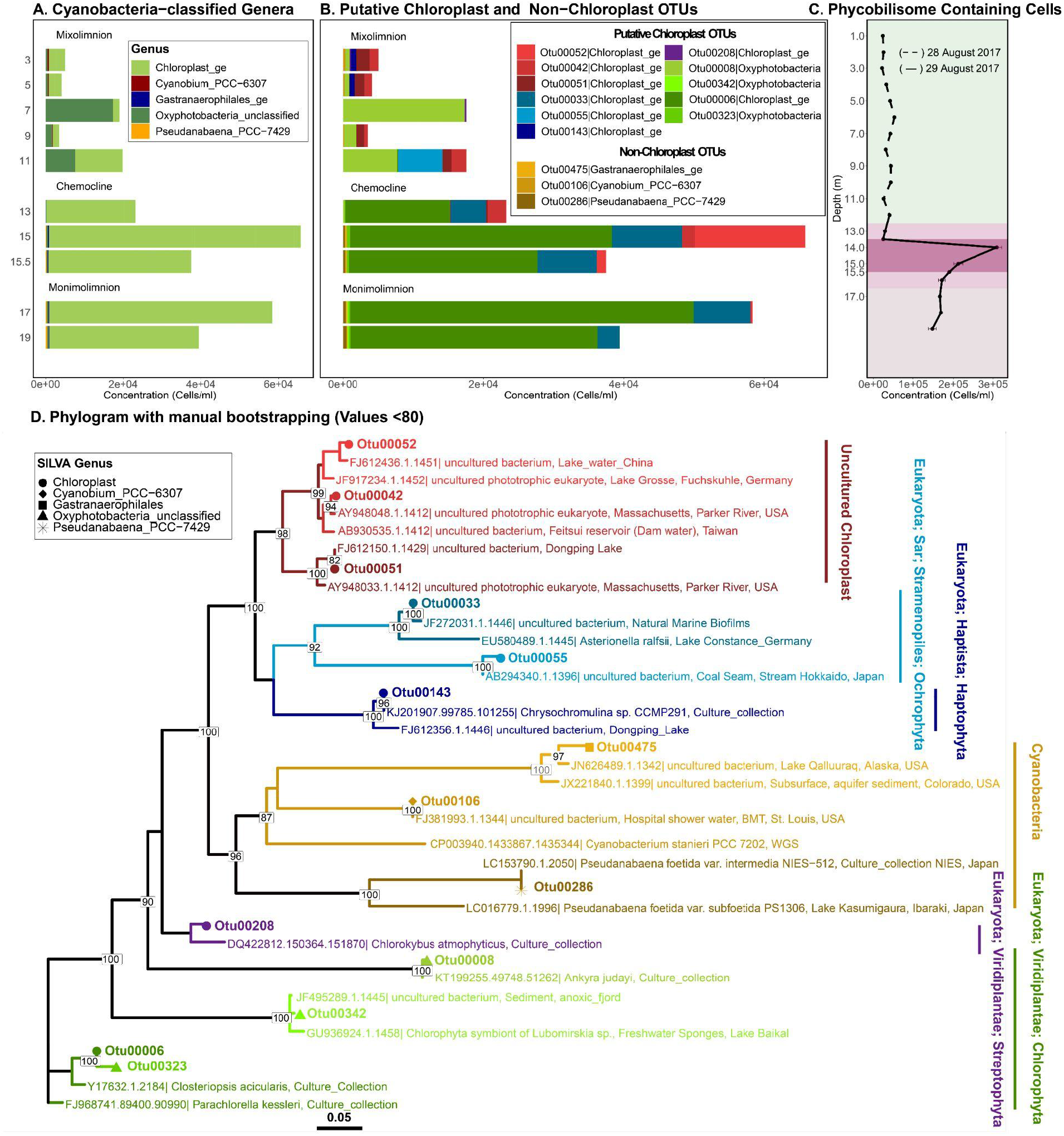
Abundances and phylogenies of Lake Cadagno phototrophs based on 16S rRNA gene amplicon sequencing and flow-cytometry. Mixolimnion samples (0-11.0 m) were collected on day 1, chemocline and monimolimnion (12.0-19.0 m) samples were collected on day 2. Abundances in panels B-C are acquired by scaling 16S rRNA gene read counts by FCM cell counts. (A) Absolute concentrations (cells/ml) of OTUs >100 cells/ml assigned to the phylum Cyanobacteria, and of those, the (B) absolute concentrations (cells/ml) of OTUs classified as putative chloroplasts and non-chloroplast at the genus level. (C) Depth profile of phycobilins-containing cell counts based on flow cytometry with a 640 nm laser. The mixolimnion, chemocline transition zone, chemocline, and monimolimnion layers are indicated by light green, lilac, purple and light brown backgrounds, respectively. (D) Phylogenetic tree of the representative 16S rRNA gene amplicon sequences of putative chloroplast and non-chloroplast OTUs in panel B, along with their two nearest neighbour sequences from the SILVA database. Clades are colour-coded consistent with the colouring of bar plots in prior panels.

In the lower mixolimnion between 7.0-11.0 m, where high oxygen, Chl *a*, and NPP persisted and peak Fv/Fm was observed (Fig. 1D, F), Otu00008, which is most closely related to the chloroplast of cultured Chlorophyta *Ankyra judayi*, dominated the subset of Cyanobacteria-classified OTUs where they represented up to 89.97% of the total cyanobacterial OTUs (Fig. 4B, D). At 11.0 m, the next most abundant OTUs were Otu00042 and Otu00055, which belonged to a clade of uncultured chloroplasts, representing 21.53% and 31.63% of the total cyanobacterial OTUs, respectively.

A primary chlorophyll peak was identified in the chemocline between 13.0-14.0 m, where FCM-based counts of phycobilin-containing cells (309,333 cells/ml; Fig.4C), phycocyanin pigments (26 µg/L day 2, Fig1D), and Chl *a* pigments (45 µg/L day 2) rose sharply (day 2; Fig. 1D). While Cyanobacteria-classified OTUs represented nearly a quarter of all OTUs at the primary chemocline chlorophyll peak, they represented only 7.28% of the total OTUs at the secondary chemocline chlorophyll peak (15 m; Table 3). The most abundant chloroplast OTU throughout the chemocline, Otu00006, was most closely related to the chloroplasts of cultured Chlorophyta, *Parachlorella kessleri* and *Closteriopsis acicularis*, a genus^33^ observed in meromictic Lake Tanganyika (Fig. 4D). The next most dominant chloroplast-like OTUs, Otu00033 and Otu00052, clustered with cultured Ochrophyta chloroplasts and uncultured chloroplasts, respectively (Fig. 4D). In the monimolimnion, Otu00006 and Otu00033 dominated the Cyanobacteria-classified OTUs (Fig. 4D), which were most closely related to Chlorophyta and Ochrophyta chloroplasts, respectively.

### Prokaryotic genotypic and phenotypic diversity trends

Bacterial genotypic diversity (16S rRNA gene) and PLP phenotypic diversity (based on individual cell features distinguished by FCM^34^) both dropped at 7.0 m and rose in the lower mixolimnion where oxygenic primary production peaked (Fig. 5A-B, Fig. 1F). Overall, genotypic alpha diversity (Shannon) trends were near-uniform, except at the peak of turbidity (15.0 m) where PLP phenotypic alpha diversity peaked and moderate genotypic alpha diversity was observed (Fig. 5A-B). Below the chemocline, consistently high genotypic and phenotypic alpha diversity was observed in the anoxic monimolimnion (Fig. 5). In the Principal Coordinate Analysis (PCoA) ordinations of bacterial community dissimilarity (Bray Curtis), the first two components combined accounted for 77% and 46.2% of all observed variation in genotypic and phenotypic beta diversity, respectively (Fig. 5 C-D). When both genotypic and phenotypic diversity was considered, the oxic mixolimnion and the co-clustering anoxic chemocline and monimolimnion samples separated along the first axis (Fig. 5 C-D). In the phenotypic diversity ordination (Fig. 5D), the mixolimnion samples separated along the second axis by whether they originated from the high-oxygen (1.0-7.0 m, light green) or the mid-oxygen (8.0-11.0 m, dark green) zones (Fig. 5 C-D). As with phenotypic alpha diversity, oxygen and light were negatively correlated with phenotypic and genotypic beta diversity (Fig. S7).

**Figure. 5:**
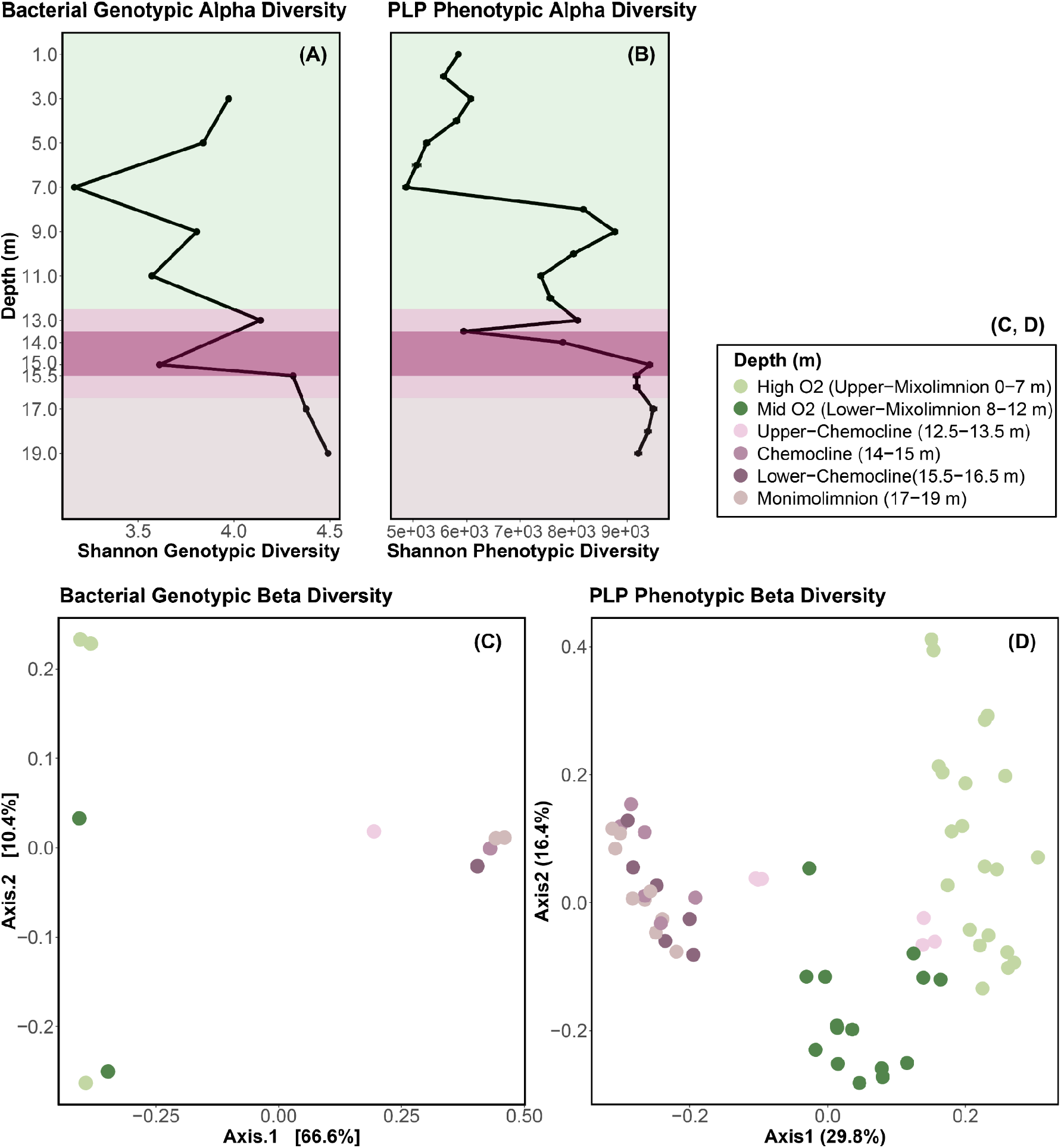
Trends in microbial community genotypic (16S rRNA gene amplicon sequencing, including bacterial and chloroplast-related OTUs) and phenotypic (PLP features based on flow-cytometry^34^) alpha and beta diversity in Lake Cadagno. (A) Variation in genotypic alpha diversity (Shannon) with depth. (B) Variation in PLP phenotypic alpha diversity (Shannon). Mixolimnion samples (0-11.0 m) were collected on day 1, chemocline and monimolimnion (12.0-19.0 m) samples were collected on day 2. Colours of data points represent depth strata sampled. (C) PCoA representing genotypic beta diversity dissimilarity (Bray Curtis) of microbial communities. (D) PCoA representing phenotypic beta diversity dissimilarity (Bray Curtis distance) of microbial communities.

## DISCUSSION

In this study, the investigation of viral, microbial and biogeochemical dynamics through Lake Cadagno’s water column improves our understanding of how biological and geochemical connections between the spatially segregated food webs in the Proterozoic ocean model^3^ may have manifested.

### Eukaryotic algae associated with oxygenic phototrophy in the mixolimnion

With the major transition to an oxygenated ocean 800-700 Ma, the ever-reducing levels of toxic sulfide and relief of nitrogen limitations likely facilitated the evolution of modern eukaryote precursors, as proposed in the Proterozoic ocean model^3^. In the oxic mixolimnion of Lake Cadagno, we observed low levels of sulfur and ammonia in combination with high abundances of eukaryotic algal chloroplast OTUs and photosynthetic activity, simultaneously with the low abundances of small (<40 µm) prokaryotic-like particles. These patterns reflect the conditions predicted during the early period of eukaryotic evolution in the Proterozoic ocean (Fig. 6A). The chloroplast sequences identified in the mixolimnion were dominated by those of the freshwater chlorophyte, *Ankyra judayi* (Chlorophyta; OTU00008; Fig. 6B), which has not yet been described in Lake Cadagno. Likely due to its ability to endure high UV radiation, *Ankyra judayi* has been observed to outcompete other phytoplankton and grow to high densities in a high-alpine Andean lake^35^, which may explain its presence in the sun-lit mixolimnion of high-alpine Lake Cadagno and may reflect traits of early photosynthesizing eukaryotes. Our findings are consistent with prior microscopy-based studies that have reported that the majority of the algal biomass was attributed to eukaryotic algae, such as *Echinocoleum* and *Cryptomonas* (Chlorophyta) and diatoms^18^. These observations of the Lake Cadagno mixolimnion support the proposed evolution of eukaryotes in the oxic strata of the Proterozoic ocean, where, though spatially segregated, they co-occurred with abundant and productive chemocline and monimolimnion bacteria in a unified ecosystem.

**Figure. 6:**
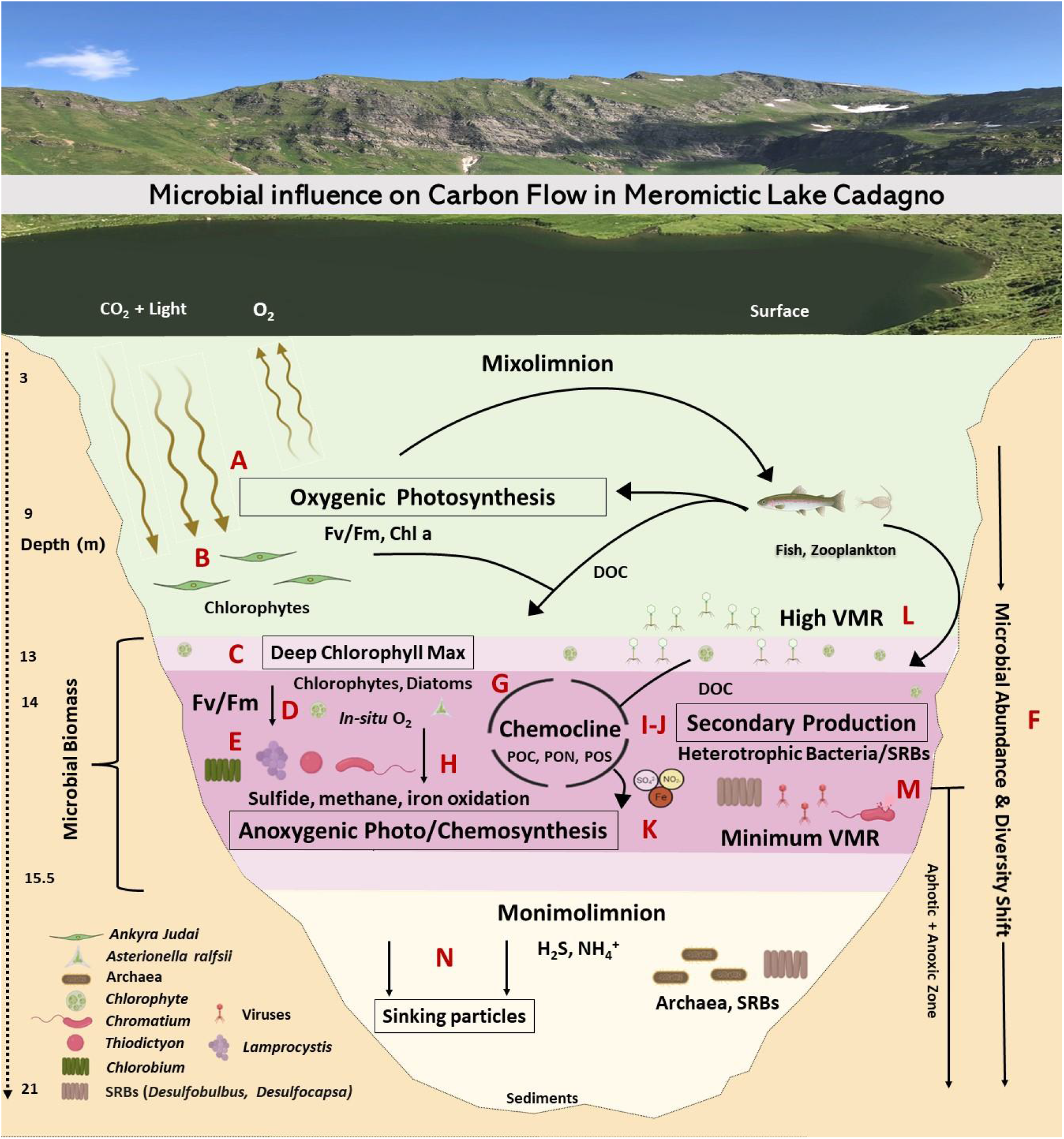
Microbial loop of meromictic Lake Cadagno.

### Coexistence of oxygenic and anoxygenic primary producers in the chemocline

The processes of the major microbial oxygenic and anoxygenic guilds have been reported to contribute almost equally to organic matter production in the Lake Cadagno chemocline ^18^, as they may have in ancient ocean chemoclines. Oxygenic (chlorophytes, diatoms, cyanobacteria) and anoxygenic phototrophs (PSB and GSB) coexisted in the Lake Cadagno chemocline (Fig. 6 C-E). Previous studies reported a cyanobacterial bloom (28 August, 12 September 2017) at the oxic-anoxic transition directly above the phototrophic sulfur bacteria ^36,37^; however, cyanobacteria (*Cyanobium, Gastranaerophilales, Pseudanabaena*) were rare in this study. Furthermore, the shift of oxygenic to anoxygenic photosynthesis is coincident with a decreasing efficiency of oxygenic photosynthetic microbes (F_v_/F_m_), increasing concentrations of hydrogen sulfide, and a shift in bacterial abundance and beta-diversity (Fig. 6F). When considering oxygenic and anoxygenic photosynthesis, the nitrogen cycle likely mediated by N_2_ fixing cyanobacteria and purple sulfur bacteria has been proposed to provide inorganic nitrogen essential for ancient ocean photoautotrophy^3,10^. The peak in PON, and the rise in biologically available N (ammonia) at the peak turbidity, suggests active N-fixation was occurring in the Lake Cadagno chemocline. N-fixation is suspected to be performed by the major carbon assimilators, PSB (*Chromatium, Thiodictyon*) and GSB (*Chlorobium*) of the chemocline, which is known to encode genes involved in nitrogen fixation^10,13,38,39^. Their diazotrophic metabolism may provide PSB and GSB the nitrogen needed for organic matter production (POC, PON, POS) in the Lake Cadagno chemocline (Fig. 6G). Such relief from nitrogen limitation may have supported the persistence of PSB and GSB in the 1.6 Gyr old ocean basin of the permanently stratified Paleoproterozoic sea^40^.

Apart from oxygenic and anoxygenic primary production, aerobic respiration has also been proposed to contribute to organic matter production in the ancient ocean chemocline ^3^. However, knowledge of which microbes are associated with such activity was missing from the Proterozoic conceptual model. *Chromatium*, identified as the most abundant chemocline microbe in this study, was previously reported to carry out aerobic sulfide oxidation that contributed up to 40% of total dark carbon fixation^41^ (Fig. 6L). This aerobic respiration by *Chromatium okenii* was coupled with *in-situ* production of oxygen by photosynthetic algae^41^, methane oxidation by methanotrophs and iron oxidation, likely by *Chlorobium* and *Rhodobacter*, where iron oxidation accounted for 10% of total primary production in the chemocline^42,43^ (Fig. 6H). Through their oxygen production, the oxygenic phototrophs (chlorophytes, diatoms, cyanobacteria) observed in the chemocline may play a critical role by enabling these oxidative metabolic processes central to Lake Cadagno’s biogeochemical cycling. These observations support the proposal of the Proterozoic ocean model of a mixed community of oxygenic and anoxygenic primary producers in the chemocline^3^. Future work targeting the ecological and metabolic interactions between chlorophytes, diatoms, cyanobacteria and chemocline bacteria at the oxic-anoxic interface of Lake Cadagno will improve our understanding of the evolution of life in ancient oceans.

### Nutrient remineralization through secondary production contributes to chemocline biomass

The purple sulfur bacteria (PSB), such as *Chromatium, Lamprocystis and Thiodictyon*, and green sulfur bacteria (GSB), such as *Chlorobium*, that dominated the Lake Cadagno chemocline, are known to be largely responsible for anoxygenic photo-chemosynthesis in this stratum (Fig. 6E) ^12,44^, In this first report of secondary production in Lake Cadagno, we found that rates were the highest in the chemocline, like primary production ^12,14,17,18^, exceeding rates in the mixolimnion and monimolimnion by 42.4- and 1.9-fold, respectively (Fig. 6I).

*Lentimicrobium*, the second most abundant genus in this report, has strongly implicated fermentation characteristics under limited light and oxygen in previous studies ^45^. Heterotrophic sulfur-reducing bacteria (SRBs, Fig. 6J), such as the observed *Desulfocapsa* and *Desulfobulbus*, are known to associate with phototrophic PSB (e.g., *Thiodictyon syntrophicum, Lamprocystis purpurea*) and were both also observed in the chemocline ^46,47^. Such heterotrophic microbes are likely involved in the decomposition of particulate organic matter (POC, PON, POS) and the supply of inorganic nutrients (iron, sulfate, and nitrogen) necessary to carry out primary production by and sustaining populations of oxygenic and anoxygenic photoautotrophs in the chemocline (Fig. 6K). These autotrophic and heterotrophic relations and the secondary productivity rates provide the evidence on microbes involved in the recycling of organic matter proposed in the Proterozoic ocean model ^3^.

### Potential for viral modulation of phytoplankton and PSB activity

Viruses have been proposed to have impacted ancient cyanobacterial mats by encoding the *nblA* gene responsible for degrading phycobilisomes, a pigment central to photosynthesis ^48^. They can also act as top-down controls on host populations and reprogram host metabolism during infection ^21,22,49^. VMR peaked in the lower mixolimnion and mixolimnion-chemocline transition where high VLP and chlorophyll concentrations and high photosynthetic activity were observed together with low bacterial abundances. Such coexistence between high viral loads and highly active phytoplankton populations could be explained by ongoing lytic viral infections of phytoplankton near the DCM (Fig. 6L). Such a scenario was observed in the phytoplankton *Ostreococcus* virus-host system, where host densities were maintained concurrently with continual lysis due to phase switching between resistant and susceptible host, a strategy supported by mathematical modelling ^50^. If this scenario is occurring, the sustained lysis of phytoplankton has the potential to release DOC in a viral shunt near the mixolimnion-chemocline transition. Such a DOC peak was observed slightly below this depth in Lake Cadagno. However, given that VMR is a feature that emerges from a number of underlying virus-microbe, viral life-history traits, and environmental co-factors ^51^, the prediction of sustained phytoplankton viral predation at the mixolimnion-chemocline transition requires empirical genomic confirmation.

Previous studies have identified genomic evidence for viral infection of the presumed major carbon assimilators in the chemocline, *Thiodictyon syntrophicum*, and *Chromatium okenii* ^12^. Genomes of these organisms sequenced from Lake Cadagno contain CRISPR elements ^38,39^, which derive from a type of acquired immunity against invading viral and plasmid DNA. This suggests viruses may play a role in the modulation of carbon, and sulfur cycles, as has been proposed in the study of a hydrothermal vent ^52^ and wetland ^53^ microbial communities, where viruses have been found to carry genes central to methanogenesis (*mcrA*) and sulfur reduction (*dsrA, dsrD*). Active viral lysis also provides a new possible mechanism to explain the previously observed low abundance of *Chromatium okenii*, which was attributed to microbial predation ^17^. Despite genomic evidence suggesting infection of these populations, VLP counts did not rise with PLP counts in the chemocline. Yet, our observation of minimum VMR (Fig. 6M) and peak microbial abundances in the Lake Cadagno chemocline support an emerging trend in stratified meromictic lakes, as the same observation was made in meromictic Ace Lake (Vestfold Hills of East Antarctica) ^25^. Such coincident low VMR and high microbial abundance have been recognized as a hallmark of the power-law relationship proposed to describe the VMR dynamics in aquatic systems ^51^. In the end, we suggest viruses as an important part of the Proterozoic ocean microbial community and to resolve the rates and roles of infection on the evolution and function of microbes that underlie Lake Cadagno biogeochemistry, especially in the chemocline, population-specific studies of viral infection are needed to explain their detailed role from ancient ocean perspectives.

Notably, the organic matter facilitated by eukaryotes, bacteria and viruses in the chemocline will be subjected to zooplankton and fish predation and the latter biomass will also eventually sink to the sediments contributing to the organic matter burial (Fig. 6N). Our empirical observations support the proposal that the chemocline (oxic-anoxic boundary) in the ancient ocean ^3^ played a crucial role in transitioning from anoxygenic to oxygenic microbial life, which dominated nutrient biogeochemical cycling in modern oceans ^54,55^.

## Conclusion

This work summarizes how biogeochemical exchange within microbial guilds may be driving biomass accumulation in the permanently stratified Lake Cadagno. The observed trends emphasized the ecological importance of microbial guilds in stratified ancient oceans. In recent years, the pangenomes of the dominant chemocline anoxygenic phototrophs (*Thiodictyon syntrophicum, Chromatium okenii*) have been explored ^38,44^. However, the genomes and metabolic potential of heterotrophic bacteria (e.g., *Lentimicrobium*) that recycle chemocline biomass in the presence of sulfur and ammonia are yet to be explored. Furthermore, eukaryotic phytoplankton in the deep chlorophyll max has been associated with the chemocline for at least two decades ^18^ and have survived hypoxia. Yet, insights into their cellular machinery that allows them to photosynthesize efficiently under limited light and oxygen are still missing. While primary producers (phytoplankton, and phototrophic sulfur bacteria) and heterotrophic bacteria are central to the realized function of the lake biogeochemistry and ecology, viruses may shuffle the repertoire of genes controlling both microbial guilds, hence modulating biogeochemical (carbon, nitrogen and sulfur) cycles of the lake. Future work will build from these observations to confirm such hypotheses.

## Methodology

### Water-sample collection and Physicochemical profiling

This study was conducted in Lake Cadagno (21 m deep), a high alpine meromictic lake situated at 1921 m above sea level in the Southern Alps of Switzerland ^56^. Water sampling and characterization of physical (turbidity, temperature, oxygen, conductivity), biological (Chl *a*, phycocyanin) and chemical (H_2_S, NH_4_^+^) parameters of the water column were performed as previously described ^16^. Water was sampled on the 28 (Day 1) and 29 (Day 2) August 2017 using a double diaphragm teflon pump (Almatech PSG Germany Gmbh) connected to acid-washed low-density polyethylene (LDPE) line deployed from a platform that was anchored above the deepest part of the Lake Cadagno (46.55087° N, 8.71152° E). The oxic stratum of the lake (0-10 m) was sampled on 28 August (Day 1), and the chemocline and anoxic part strata (11-19 m) were sampled on 29 August 2017 (Day 2).

Vertical lake profiles were determined using an autonomous conductivity temperature depth sensor (CTD; Ocean Seven 316 Plus CTD; IDRONAUT, S.R.L.). The CTD recorded pressure (dbar), temperature (°C), conductivity (mS/cm), and dissolved oxygen (mg/L) were measured with a pressure-compensated polarographic sensor. The error in oxygen profiles for day two was manually adjusted by referring to day 1 measurements which indicated approximately zero oxygen within and below chemocline. CTD profiles from the whole water column (0-20 m, Day 1 and 2) were used to identify the distribution of the mixolimnion, chemocline and monimolimnion and guided water sampling strategy. The CTD was equipped with an LI-192 Underwater Quantum Sensor (Li-Cor Biosciences; NE, USA) that continuously recorded photosynthetically active radiation (PAR-w/m^2^;) and a TriLux multi-parameter algae sensor (Chelsea Technologies Ltd; Surrey, UK) that measured in-vivo Chl *a* (Chl *a*, µg/L), an indicator of phytoplankton (including eukaryotic autotrophs and cyanobacteria) biomass ^57^, and phycocyanin (µg/L), a pigment characteristic of cyanobacteria ^58^.

### Chemical parameters

#### 2.1 Dissolved compounds

Samples for the analysis of dissolved compounds (DOC) were filtered through an Acropack filter cartridge (PALL) with a 0.8 µm pre-filter and 0.2 µm final filter. Filters were collected in 30 mL acid-washed and pyrolyzed pyrex tubes filled to the top and supplemented with 100 µL HCl 2M. Tubes were stored in the dark at 4° C until processed (November 2017; University of the Geneva) using a Shimadzu TOC-LCPH analysis system. Blanks consisting of Milli-Q water were made and calibration was done using the standard from 0.2 to 5 ppm. Analyses of dissolved ammonium (MNH_4_^+^) and hydrogen sulfide (H_2_S) were done on-site at the Alpine Biology Centre (CBA; Piora, Switzerland) with freshly collected water samples using a UV-visible spectrophotometer (DR 3800, HACH) ^59^.

#### 2.2 Particulate compounds

Particulate organic carbon (POC), nitrogen (PON) and sulfur (POS) were collected by filtering 150 to 500 mL of lake water (volume necessary to begin to clog the filter) on pre-combusted 47 mm GF/F filters (5h at 550 °C, cat. #WH1825-047, Whatman, Wisconsin, USA), which were then acidified (500 µL HCL 1 M, added twice at 1 h interval) and dried in an oven (65 °C) overnight in acid-washed pyrex petri dishes. Samples were sealed using parafilm and aluminium foil until analysis (November 2017, University of Geneva) using an Elemental Analyzer (2400 series II CHNS/O Elemental Analysis, PerkinElmer). Procedural blanks using MilliQ water were performed (24-25 November 2017) in duplicate to measure the background signal, which was subtracted to sample analysis. POC, PON, POS were expressed in mg/L (ppm).

### Biological parameters

#### 3.1 Primary production

Primary productivity was determined using incubation with NaH_14_CO_3_ (Perkin Elmer cat. #NEC086H005MC; Waltham, MA, USA; 5 mCi, 1 mL in glass ampoule) that was freshly diluted into 4 mL MilliQ water at pH 9.6 (adjusted using NaOH, Sigma) and added at a dose of 1 mCi/L of lake water, as described ^60^. Incubation was carried out along an incubation chain deployed at six depths for 23 h on 28-29 August 2017. Each depth consisted of a metallic ring fixed to a rope at the desired depth (3, 5, 7, 9, 11 and 13 m). The ring was surrounded by six arms on a horizontal plane, each of which ended with a bottle holder. 75 mL acid-washed glass bottles were filled to the top with lake water (ca. 100 mL) at the corresponding depth and spiked with ^14^C prior to being deployed on each level of the incubation chain. At each depth, three transparent bottles were incubated for activity measurements at *in situ* natural light intensities, and three amber bottles were incubated for dark community respiration measurements. At the end of the incubation, each bottle was filtered onto 47 mm GF/F filters (cat. #WH1825-047, Whatman, Wisconsin, USA) that were then each stored in a plastic petri dish. Petri dishes were placed in a sealed box containing calcium hydroxide (slaked lime) powder. After adding HCL (1M, 500 µL) onto filters, inorganic carbon was allowed to degas overnight in a sealed box. Filters were then collected into 20 ml scintillation vials, supplemented with 10 mL of Ultima Gold AB cocktail, and shaken manually. Slaked lime, Decon solution and pipet tips were collected as radioactive waste for appropriate disposal at the University of Geneva, while all other labware was decontaminated by using warm (40 to 50° C) 5% solution of Decon 90 (cat no. #10335650, East Sussex, England). Primary productivity was expressed in µmol of carbon net fixation per h either per L of lake water or per Chl *a*, considering a dissolved inorganic carbon concentration of 1.8 mM C ^14^.

#### 3.2 Maximum quantum yield

The maximum quantum yield (F_v_/F_m_) informs the photosynthetic health and biological activity of algae-based on photo-physiological characteristics of the photosystem II variable Chl *a* fluorescence. Maximum quantum yield was determined using a Fast Repetition Rate Fluorometer (FRRF, FastOcean PTX coupled to a FastAct base unit; Chelsea Technologies). Water for maximum quantum yield measurements was pre-concentrated 10-fold using a 47 mm filtration unit (cat. #300-4000; Nalgene), a 0.2 µm 47 mm polycarbonate filter (cat. #GVWP04700, Millipore; Darmstadt, Germany) and a 15 mL pipette to gently resuspend the cells as the sample was passed through the filter using a hand pump (pressure below 15 mbar). This step was done to ensure high enough method sensitivity and has been shown to alter neither Chl *a* nor cell integrity of samples from Lake Geneva ^61^. The FRRF was used in single turnover mode to record, in a 45 min dark-adapted natural sample, the basal (F_b_) and the maximal (F_m_) fluorescence following exposure to intense light. The sample was then gently filtered on a 0.2 µm 47 mm syringe filter (cat. #GVWP04700, Millipore; Darmstadt, Germany) to record the residual fluorescence (F_r_). For each sample, the F_b_-F_r_ represented the initial dark-adapted fluorescence F_0_ associated with intact cells. The F_v_/F_m_ was then calculated using the ratio: (F_m_- F_0_)/F_m_.

#### 3.3 Flow-cytometry (FCS)

A BD Accuri C6 cytometer (Becton Dickinson, San José CA, USA) was installed with two lasers (blue: 488nm, red: 640 nm) and four fluorescence detectors (lasers 488 nm: FL1 = 533/30, FL2 = 585/40, FL3 = 670; laser 640 nm: FL4 = 675/25) was used to estimate total cell counts from fresh unpreserved collected samples on-site in the Alpine Biology Centre. FL4 detectors were specifically used to detect emission from phycobilin, a pigment proxy for cyanobacteria. Cells were stained with SYBR green I (cat. #S7563, Molecular Probes; Eugene, OR) with a ratio of 1:10,000 (vol/vol), following incubation for 13 minutes at 37°C in the dark ^11^. Histogram of event counts versus green fluorescence (FL1 > 1,100) allowed quantification of total cells.

VLPs were counted using flow-cytometry after filtration of lake water through 55 µm mesh and 0.22 µm filters to remove large particles and cells. In triplicate, 1 ml sample from the filtrate was fixed by adding 10 µL of glutaraldehyde (25% stock solution) in sample cryovials. The samples were fixed for 15 minutes at room temperature and subsequently stored in liquid nitrogen for shipping then stored at -80°C until processed in February 2018. Virus-like particle counts were obtained using a FACSCalibur flow cytometer (Becton Dickinson Biosciences; Grenoble, France) at Université Savoie Mont-Blanc. Samples were thawed at 37°C then diluted in a sterile buffer solution of Tris-EDTA (0.1 mM Tris-HCL and 1 mM EDTA, pH 8) that had been autoclaved and filtered through 0.02 µm. Samples were then stained for 10 minutes at 75°C using SYBR Green I (Molecular Probes) at a 1:10,000 final concentration ^62^.

##### 3.3.2 Phenotypic alpha diversity of PLP (Prokaryotic-like-particles)

Raw flow-cytometry files were used to estimate phenotypic traits (morphology and nucleic acid content) using the Phenoflow package in R ^34,63^. Briefly, this approach estimates kernel densities on multiple bivariate single-cell parameter combinations (e.g., fluorescence and scatter intensity) and concatenates these into a feature vector, which can be thought of as a phenotypic fingerprint ^34^. This fingerprint represents community structure in terms of the phenotypic attributes of the cells that comprise the community—essentially, any feature that influences the way the laser interacts with passing particles, such as morphology, size, and nucleic acid content, is captured in the phenotypic fingerprint. Using this approach, bacterial phenotypic diversity metrics have been shown to be highly correlated with community taxonomic diversity^34^. To calculate phenotypic alpha diversity, phenoflow calculates Hill number diversity indices (D_q_) from order 0-2, where 0 represents the richness and q>1 represents richness and evenness, also referred to as D0 (richness), D1 (Shannon), and D2 (inverse Simpson) ^64^. Phenotypic beta diversity was estimated using PCoA of phenotypic fingerprints.

Fluorescence detectors (lasers 488 nm: FL1-H = 533/30, FL3-H = 670) were used to target PLP diversity. Polygon gating was applied to filter out the noise observed in control samples (Fig. S1). Clockwise, the polygon gate coordinates (x,y) were: (7, 6), (15, 6), (15, 17), and (7,10). Kernel density estimates were calculated by applying binning grid (128 × 128) to FSC-H (Forward scatter height), SSC-H (Side scatter height), FL1-H and FL3-H. The obtained kernel densities values were assigned into one-dimensional vectors, termed *phenotypic fingerprints*.

#### 3.4 Secondary production

Heterotrophic bacterial production, also termed secondary production, was estimated by the micro-centrifuge ^3^H-Leucine bacterial production method ^65^, as described ^66,67^. A working solution of 5 µCi mL^-1^ leucine was made from a manufacturer stock solution of radioactive leucine, L-[3,4,5-^3^H(N)] (cat. #NET460A005MC, PerkinElmer, MA, USA). To reduce the potential for autodegradation of the leucine radiolabel, this stock ^3^H-Leucine material was stored in the dark at 2 to 4° C (never frozen) and used for the incorporation experiment within 5 days. Two replicates and one trichloroacetic acid (TCA)-killed control (5% [vol/vol] final concentration; cat. 91228, Sigma) were used for each depth. Briefly, 1.5-mL of lake water samples were incubated with a mixture of radioactive and nonradioactive leucine at final concentrations of 20 nM. Using the [^3^H]-Leucine working solution, enough “hot” leucine was added to each tube to create a sufficient signal (50% or 85% of hot leucine were used in the mixture for samples collected above and below 13m depths, respectively). Incubations were conducted in a dark incubator at the respective *in situ* temperatures (corresponding to the depths) for 2 h. Saturation of leucine incorporation over this period was tested with 20, 30 and 40 nM of total leucine. At the end of the incubation, 200 µL of 50% TCA is added to all but the control tubes to terminate leucine incorporation. To facilitate the precipitation of proteins, bovine serum albumin (BSA; Sigma, 100 mgL^-1^ final concentration) was added, then samples were centrifuged at 16,000 g for 10 min ^68^. The supernatant was discarded and the resultant precipitated proteins were washed with 1.5 mL of 5% TCA by vigorous vortexing and again centrifuged (16,000 g for 10 min). The supernatant was discarded. Subsequently, 1.5 mL of UltimaGoldTM uLLt (Part number: 6013681, PerkinElmer, MA, USA) was added to each vial, mixed, and allowed to sit for >24 h before the radioactivity was determined by liquid scintillation using a Beckman LS6500 liquid scintillation counter. To calculate biomass production, the following conversion factors were applied: molecular weight of leucine of 131.2 g/mol, a fraction of leucine per protein of 0.073), the ratio of cellular carbon to the protein of 0.86, and 2.22×10^6^ DPM/μCi to convert disintegrations per minute (DPMs; the measure of radioactivity) to μCi. A factor of 1.55 kg C mol leucine^-1^ was used to convert the incorporation of leucine to carbon equivalents, assuming no isotope dilution ^69^.

#### 3.5 Statistics

The magnitude and significance of linear relationships between physical and biological parameters were determined with linear regression scatter plots in R-Studio (1.4.1106) using ggplot2 (3.3.3)^70^ and ggpubr (0.4.0) ^71^ packages which included correlation coefficients (R) with p-values to the scatter plots ^72^. R>0.70 was considered a strong correlation, and a p-value <0.05 was considered statistically significant.

### Illumina MiSeq 16S rRNA gene amplicon sequence analyses

#### 4.1 16S rRNA gene amplicon sequencing

The samples were collected from 3.0, 5.0, 7.0, 9.0, 11.0, 13.0, 15.0, 15.5, 17.0 and 19.0 m to capture Lake Cadagno’s mixolimnion (surface to 10.0 m), chemocline (13.0-15.5 m), and monimolimnion (below 16 m). Collected water samples (20 L/depth) were 55 µm pre-filtered to remove large particles and aggregates and then 0.22 µm filtered (cat. #GPWP14250 142 mm Express Plus Filter, Millipore; Darmstadt, Germany) using a peristaltic pump. Filters were flash-frozen in liquid nitrogen and stored at (−80°C) until extraction of DNA using the DNeasy Blood & Tissue Kit (cat. #69504 QIAGEN, Germantown, MD, USA) combined with QIAshredder (cat. #79654, QIAGEN, CA, USA), as described ^73^. The first and second elutions of DNA (50 µl each) were stored (4° C) until sequencing. Dual indexed primers were used and the V4 region of the 16S rRNA gene amplicon was targeted and amplified, as described ^74^. Sequencing libraries were prepared using the Illumina Nextera Flex kit (Illumina; CA, USA) and sequenced using the MiSeq platform (500 cycles, Illumina) at the University of Michigan Microbiome Sequencing Core.

As microbes can have multiple 16S rRNA gene copies per cell, the sequenced reads were quality controlled, assembled, trimmed, aligned, and clustered at 97% identity threshold into a representative operational taxonomic unit called OTU using mothur^75^. The OTUs were taxonomically classified using SILVA ^76^ and TaxAss freshwater ^77^ 16S rRNA gene databases. The code used for running the mothur analysis is available at the project GitHub repository (^78^/). Raw sequences are available under project accession ID PRJNA717659 at NCBI.

#### 4.2 Genotypic alpha and beta diversity and Phylogenetic Tree

Downstream analysis was carried out by the Phyloseq package in R-studio (v1.3.1093) to analyze the alpha and beta genotypic diversity of bacterial communities. Taxa not observed in 20% of samples at least three times were removed. The Alignment, Classification, and Tree (ACT) function of SILVA were used to generate the phylogenetic tree ^79^. Representative sequences of chloroplast OTUs were aligned to SILVA sequences (SINA 1.2.11) Two neighbours per query sequence (95% minimum sequence identity) were added to the tree with the chloroplast OTU sequences. Advance variability profile was selected as ‘auto’ and sequences below 90% identity were rejected. The phylogenetic tree of the OTUs was bootstrapped with RAxML ^80^ with 20 maximum likelihood (ML) and 100 bootstrapped searches. Annotation was performed in R-studio (1.4.1106) with package ggtree (2.2.4) and Adobe Illustrator (25.2.1).

#### 4.3 Absolute Abundances (cells/ml)

The absolute quantifications of taxa were determined as described ^31^. Briefly, the relative abundance of bacterial and archaeal OTUs of a given taxon (taxon read count/total read count in sample) was multiplied by the PLP concentrations (cells/ml) for that sample, as determined by flow cytometry (FCS). Thereon, archaeal abundance was deducted from the total prokaryotic population to get an absolute abundance of the bacterial population. Because chloroplasts likely originated from Cyanobacteria ^81^, their OTUs are assigned under phylum Cyanobacteria by SILVA.

Note: as per the previous methodological findings, both relative and absolute abundance were referred for interpretation of Lake Cadagno microbial population ^31^.

## Supplementary

[Table 1](https://docs.google.com/spreadsheets/d/1S1buluH3MS2nZFFnw1txogTBsSwrzOUyKVoCz5djW0I/edit?usp=sharing): The relative and absolute abundance of bacterial communities, including chloroplasts.

[Table 2](https://docs.google.com/spreadsheets/d/1u98I3yphaFMYHinv76aG5gfMdpP_dq7JWHUg8Cyhbd8/edit?usp=sharing): The relative and absolute abundance of archaeal communities.

[Table 3](https://docs.google.com/spreadsheets/d/1i40Y_MurPoQbyDDt27eesoiUmBf_hkHKM-seq-lZvsc/edit#gid=1551248890): The relative and absolute abundance of cyanobacterial communities, including chloroplasts.

[Data and code availability](https://github.com/DuhaimeLab/Lake_Cadagno_microbial_loop_Saini_et_al_2021)

## Acknowledgements

The project was funded by Swiss National Science Foundation grant no. PP00P2-138955. We thank the University of Michigan microbiome core facility for sequencing Lake Cadagno 16S data. Many thanks to Dr Sandro Peduzzi, and to the Alpine Biology Center Foundation (Switzerland) in general, for providing housing during Lake Cadagno sampling, and for offering general comments on the manuscript. We also acknowledge BioRender for providing microbiology icons used in Fig. 6 schematic. Lastly, I, Jaspreet, express sincere gratitude to the Swiss Confederation and Ernst and Lucy Schmidheiny Foundation for financially supporting me with a monthly salary which aided the development of this manuscript.

## Supplementary figures

**Fig. S1:**
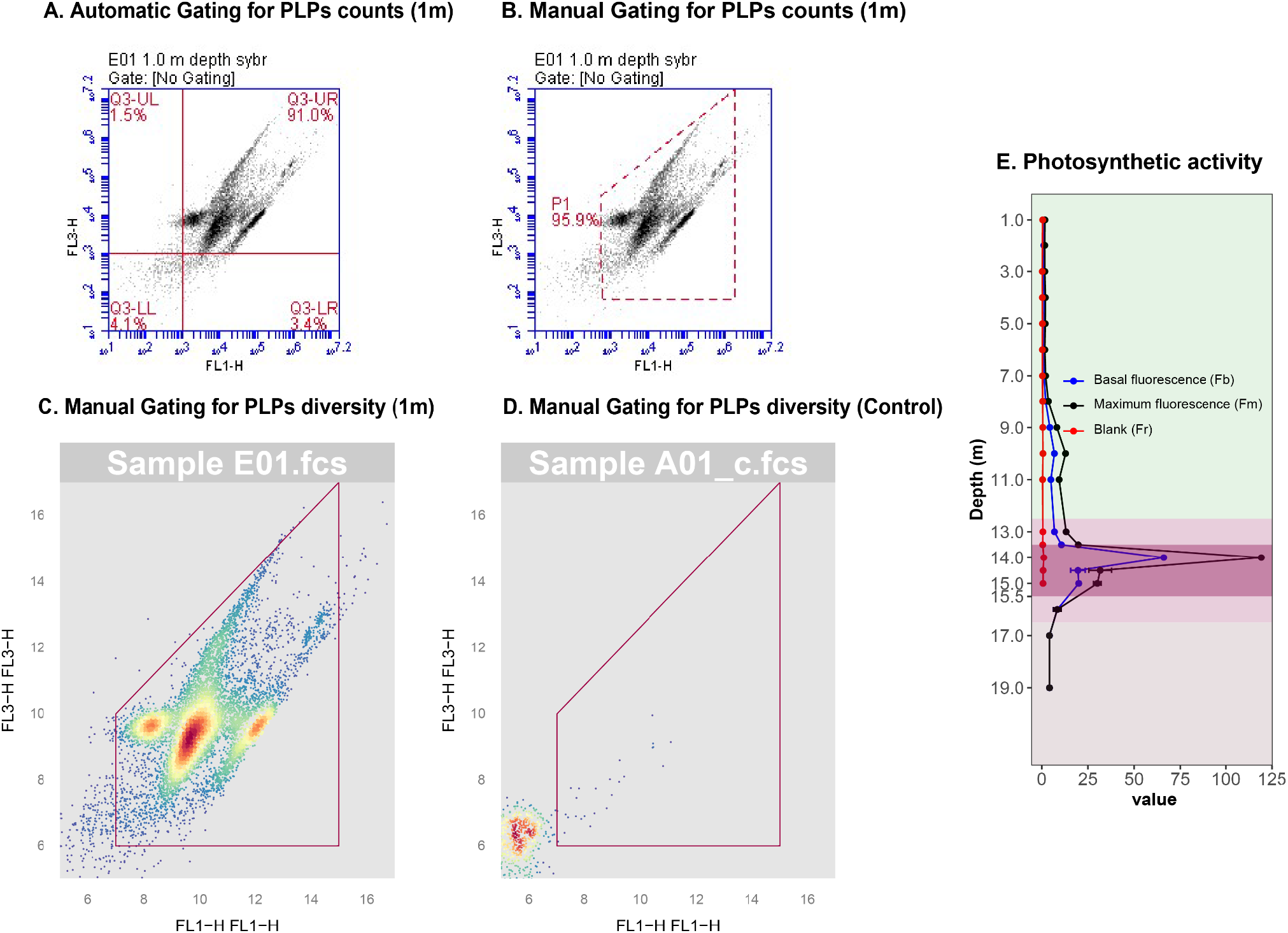
Raw flow-cytometry files visualized with; BD6 flow-cytometer interface (A-B) and R-studio Phenoflow package (A-D). The red outline represents the gating strategy for cell counting and phenotypic diversity analysis. FCM events representing prokaryotic-like particles in a Lake Cadagno (1 m depth) targeted using (A) Four-area layout vs (B, C) Manual coordinated based gating. (D) FCM events represent prokaryotic-like particles in a 0.2 μm filtered, cell-free MilliQ water (control). (E) Experiment to monitor the health of photosynthetic cells. Basal fluorescence (F_b_) and maximum fluorescence (F_m_) were obtained to calculate Fv/Fm values.

**Fig. S2.**
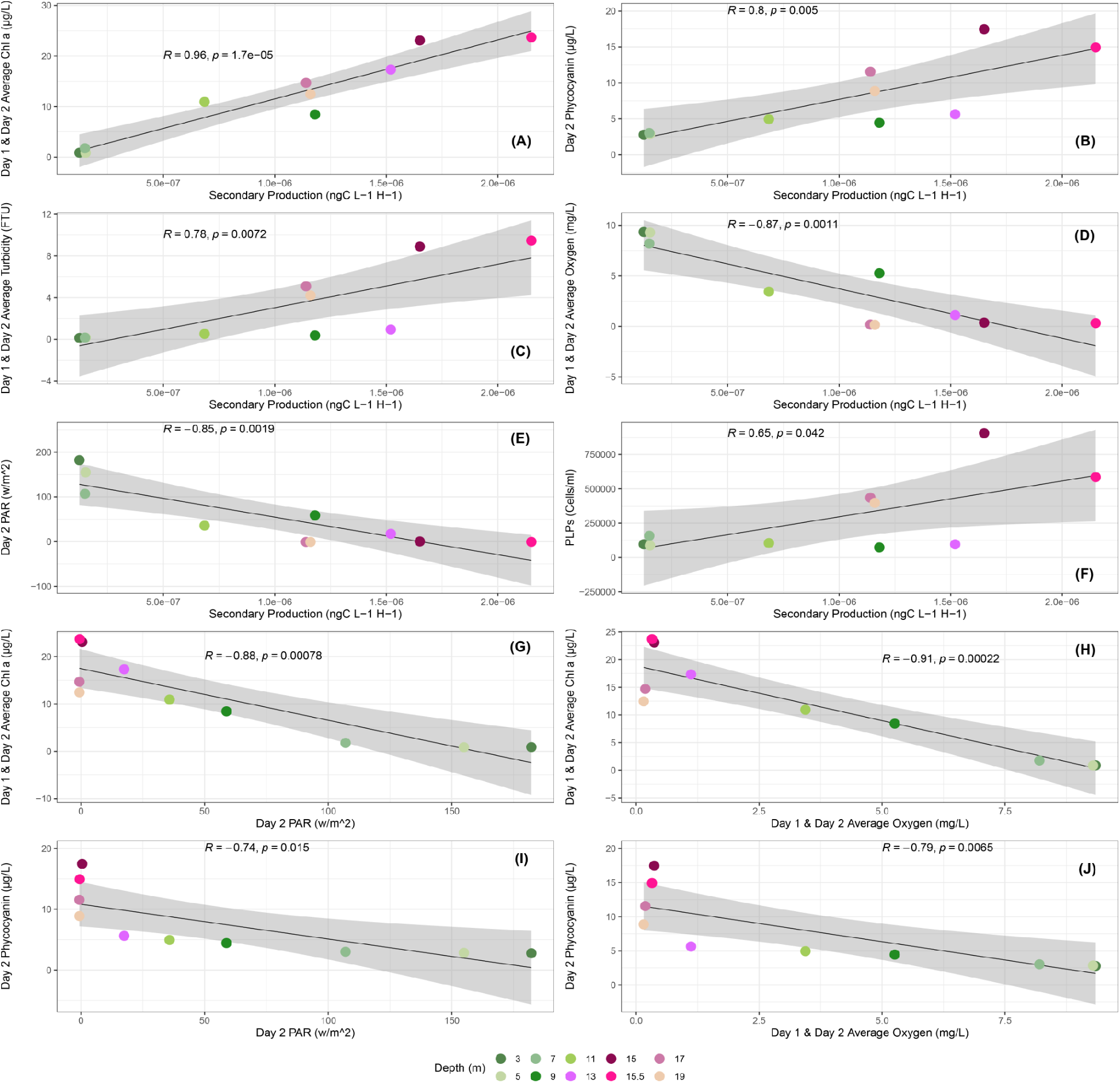
R-coefficient and p-values were obtained using linear regression performed between secondary production, photosynthetic pigments, Prokaryotic-like-particles (PLPs), and physicochemical parameters including turbidity, oxygen and light. Lake depth in meters (m) is indicated by colors. (A) Secondary production vs Chl a (B) Secondary production vs Phycocyanin (C) Secondary production vs Turbidity (D) Secondary production vs Oxygen (E) Secondary production vs light (F) Secondary production vs PLPs (G-J) Chl a and Phycocyanin vs oxygen and light.

**Fig. S3.**
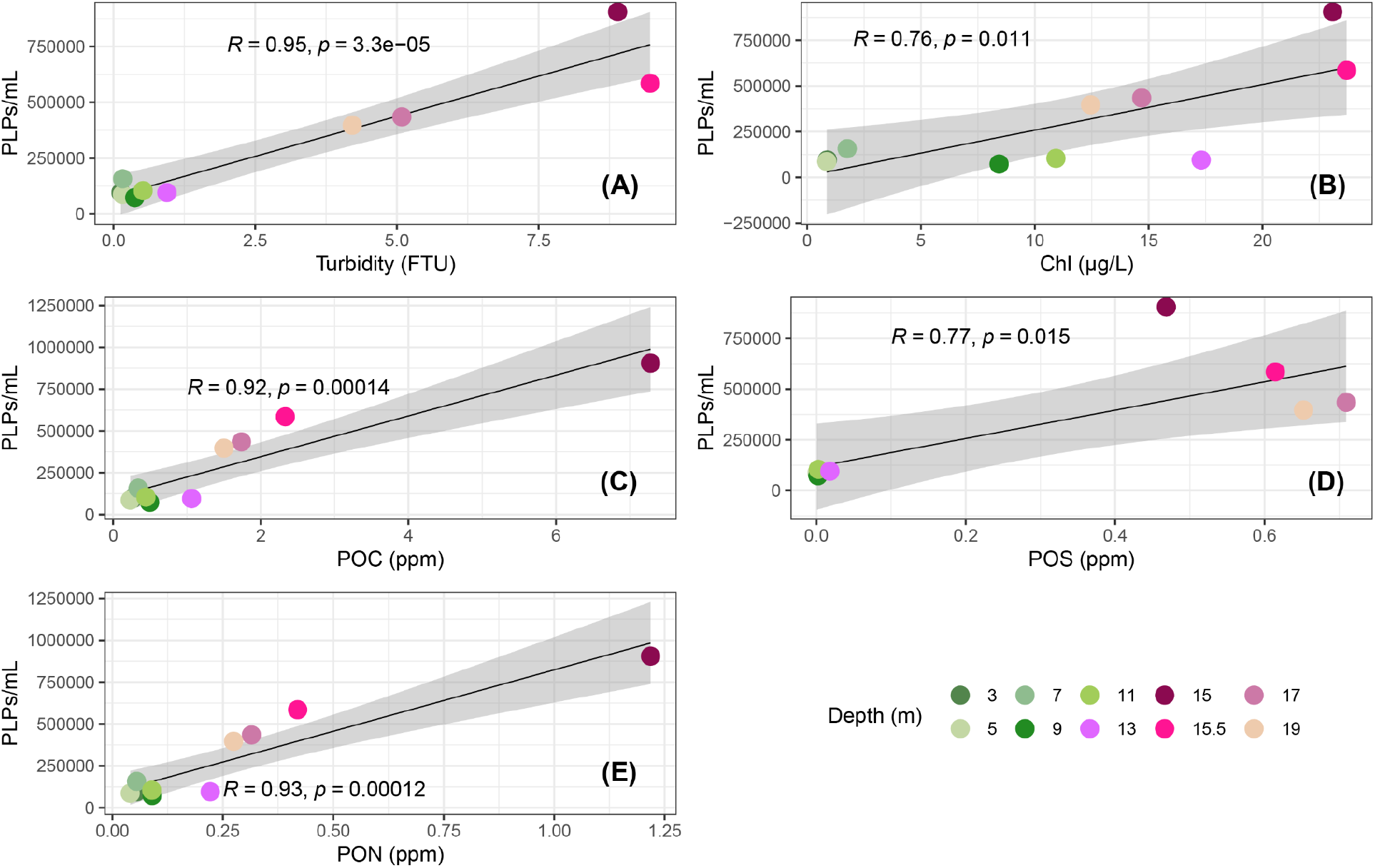
R-coefficient and p-values were obtained using linear regression performed between Prokaryotic-like-particles (PLPs) and biomass indicating factors. Lake depth in meters (m) is indicated by colors. (A) PLPs vs Turbidity (B) PLPs vs Chl a (C) PLPs vs particulate organic carbon (POC). (D) PLPs vs particulate organic sulfur (POS). (E) PLPs vs particulate organic nitrogen (PON).

**Fig. S4.**
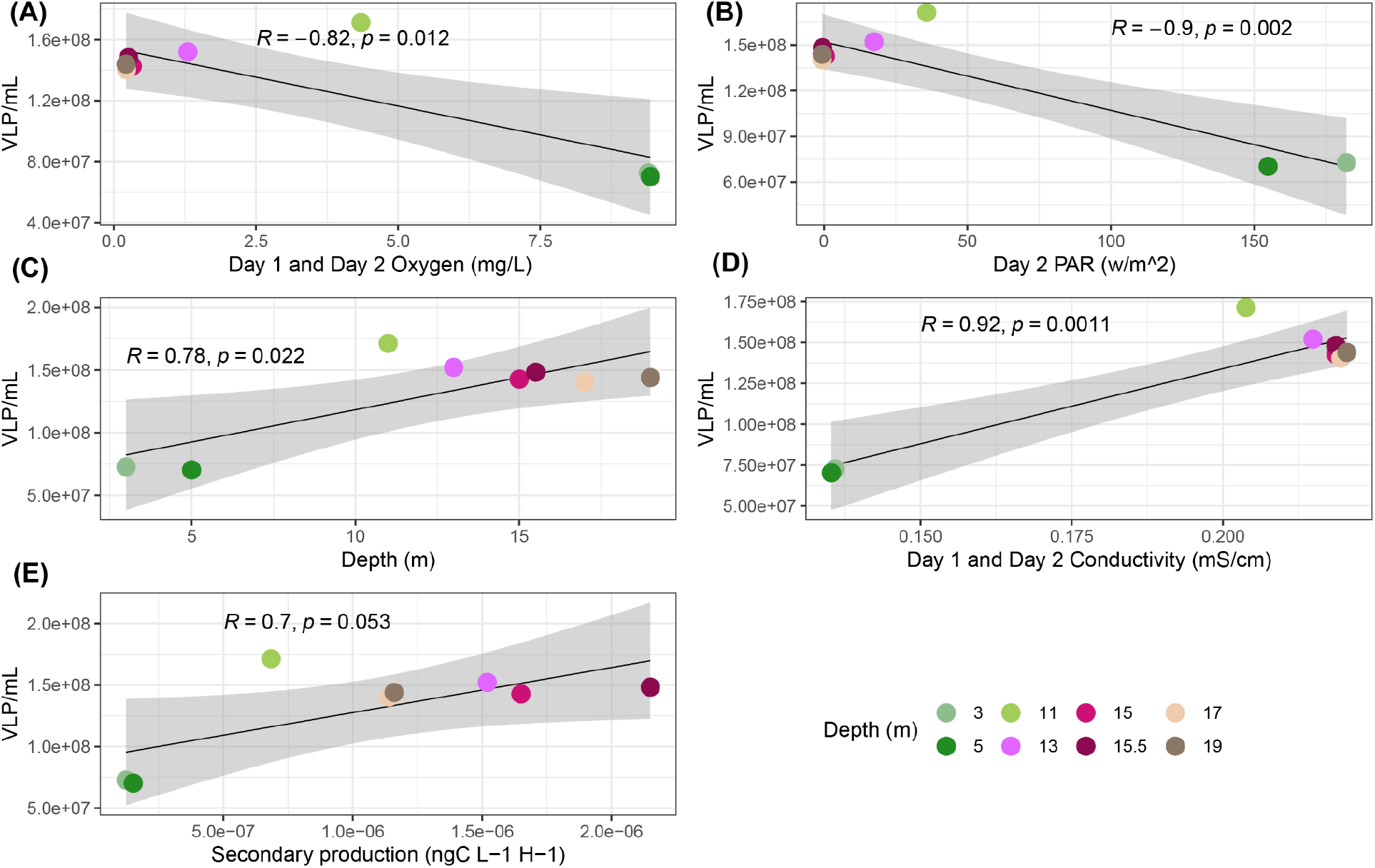
R-coefficient and p-values were obtained using linear regression performed between Virus-like-particles (PLPs) and physicochemical parameters. Lake depth in meters (m) is indicated by colors. (A) VLPs vs Oxygen (B) VLPs vs Light (C) VLPs vs depth. (D) VLPs vs conductivity. (E) VLPs vs secondary production.

**Fig. S5.**
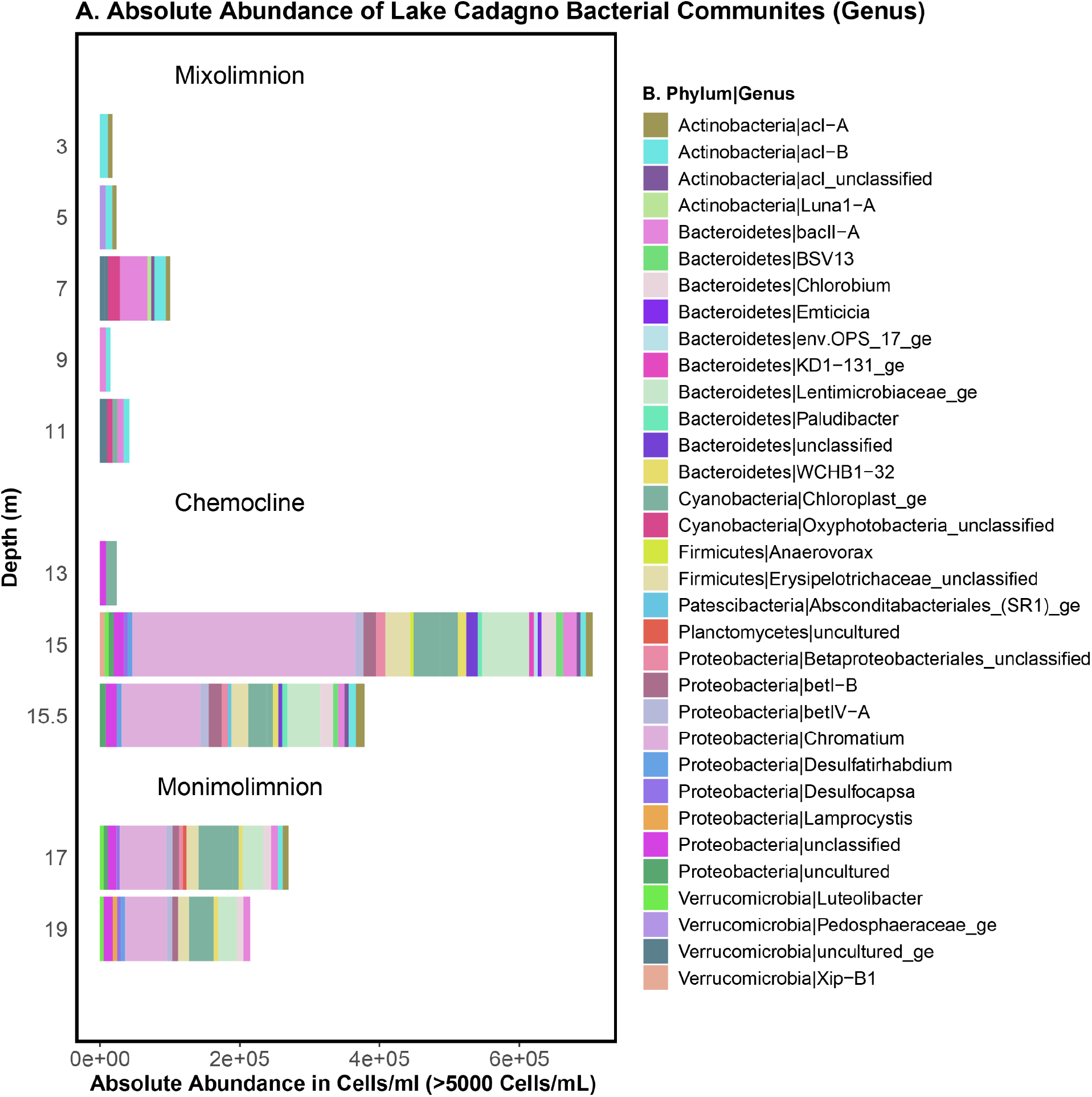
Absolute abundance of microbial communities at genus level within the vertical water column of Lake Cadagno.

**Fig. S6.**
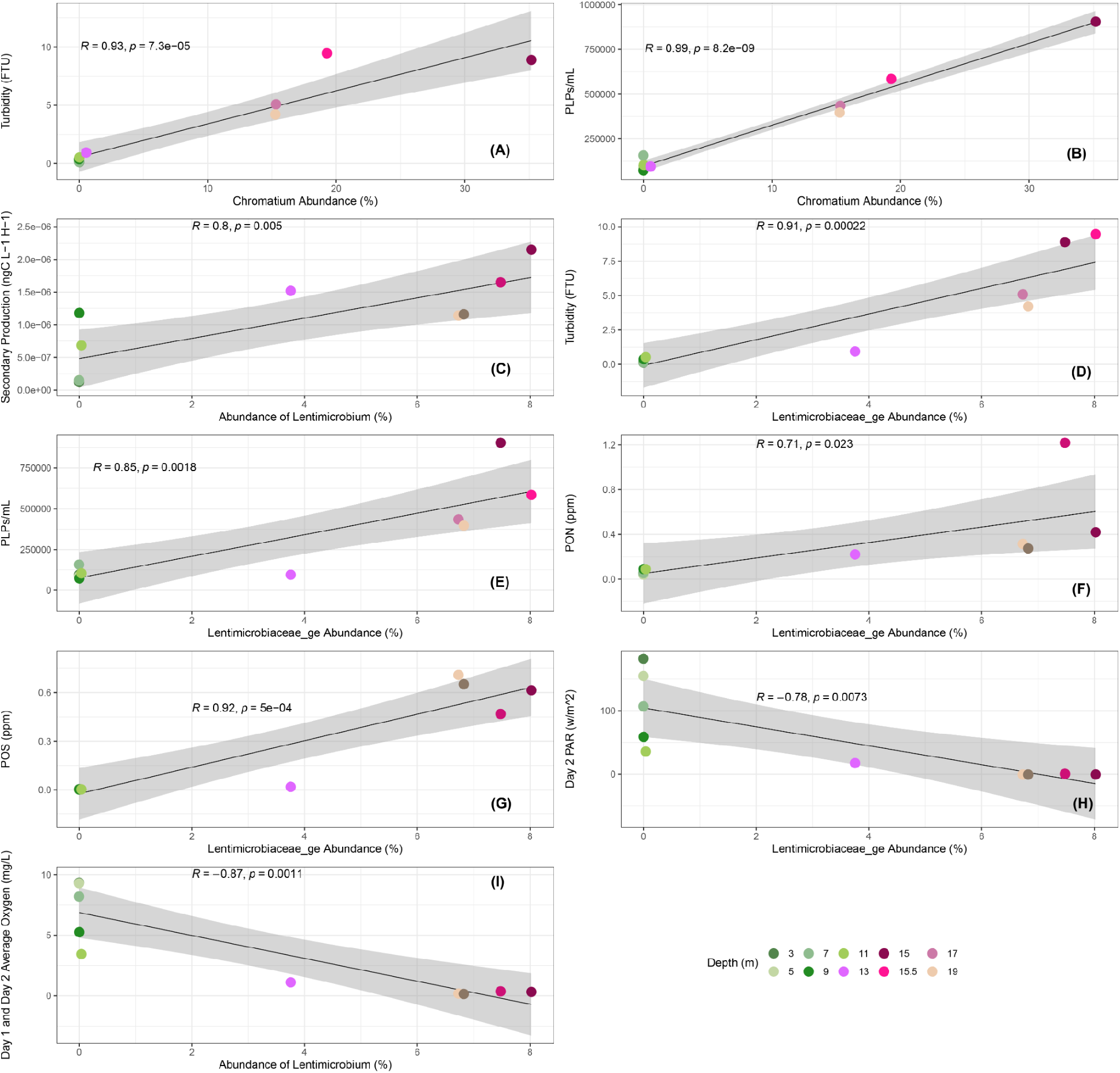
R-coefficient and p-values were obtained using linear regression performed between microbial abundance and physicochemical parameters. Lake depth in meters (m) is indicated by colors. (A-B) Chromatium vs PLPs and turbidity. (C-I) Lentimicrobium vs secondary productivity, turbidity, PLPs, PON, POS, light and oxygen.

**Fig. S7.**
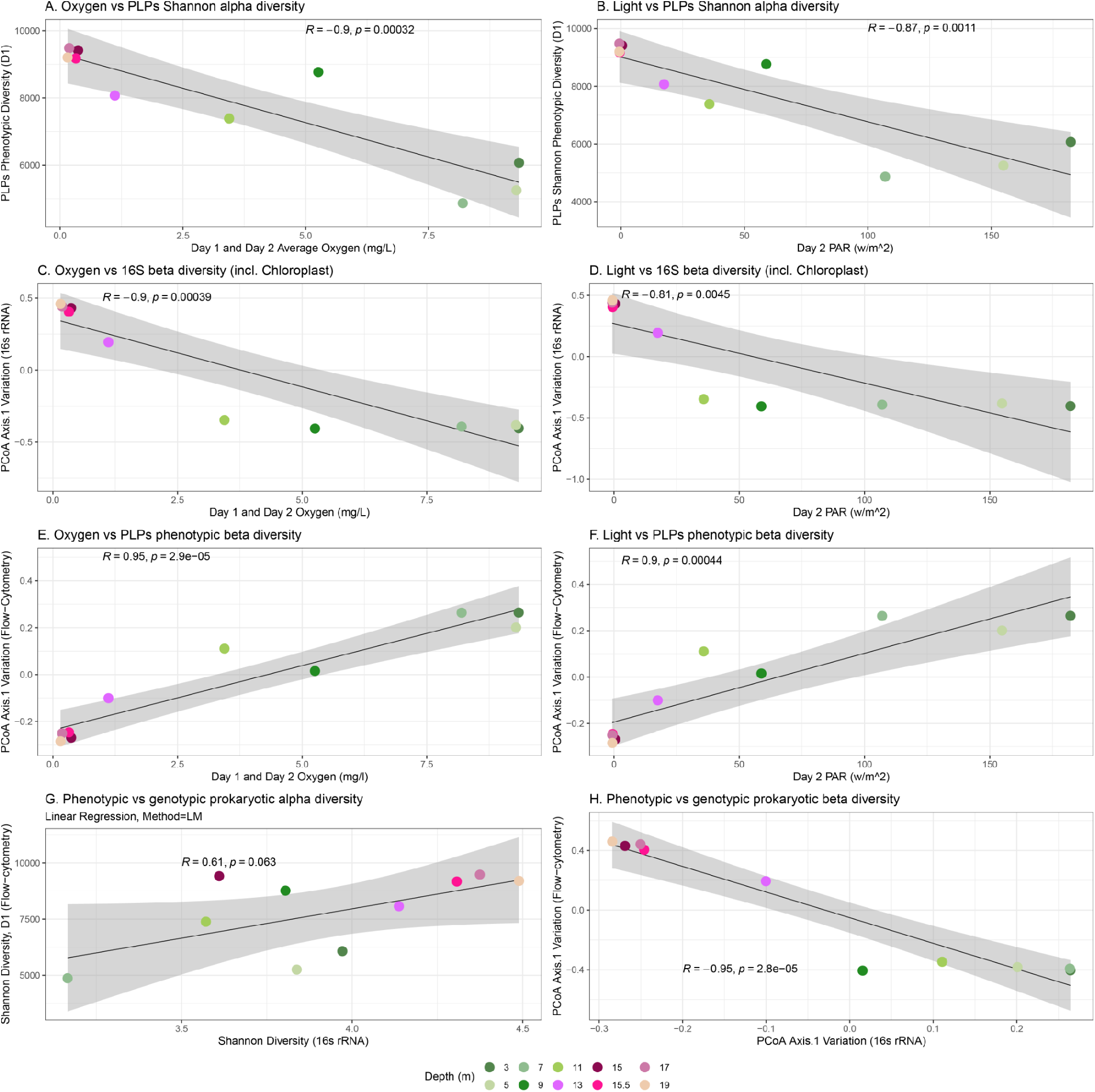
R-coefficient and p-values were obtained using linear regression performed between microbial alpha and beta diversity and physicochemical parameters. Lake depth in meters (m) is indicated by colors. (A-B) PLPs Shannon alpha diversity vs oxygen and light. (C-D) 16S beta diversity vs oxygen and light. (E, F) PLP phenotypic beta diversity vs oxygen and light. (G-H) Phenotypic vs genotypic prokaryotic alpha and beta diversity.

